# Mutanofactin promotes bacterial adhesion and biofilm formation of cariogenic *Streptococcus mutans*

**DOI:** 10.1101/2020.08.22.262196

**Authors:** Zhong-Rui Li, Yongle Du, Jin Sun, Aifei Pan, Lin Zeng, Roya Maboudian, Robert A. Burne, Pei-Yuan Qian, Wenjun Zhang

**Affiliations:** Department of Chemical and Biomolecular Engineering, University of California, Berkeley, CA, USA; Department of Ocean Science and Hong Kong Branch of the Southern Marine Science and Engineering Guangdong Laboratory (Guanzhou), The Hong Kong University of Science and Technology, Hong Kong, China; State Key Laboratory for Manufacturing Systems Engineering, Xi’an Jiaotong University, Xi’an, China; Department of Oral Biology, College of Dentistry, University of Florida, Gainesville, Florida, USA; Chan Zuckerberg Biohub, San Francisco, CA, USA

## Abstract

Cariogenic *Streptococcus mutans* is known as a predominant etiological agent of dental caries due to its exceptional capacity in forming biofilms. From strains of *S. mutans* isolated from dental plaque, we here discover a polyketide/non-ribosomal peptide biosynthetic gene cluster, *muf*, which directly correlates with a strong biofilm-forming capability. We then identify the *muf*-associated bioactive product, mutanofactin-697 that contains a novel molecular scaffold, along with its biosynthetic logic. Further mode-of-action studies reveal mutanofactin-697 binds to *S. mutans* cells nonspecifically, increases bacterial hydrophobicity, and promotes bacterial adhesion and subsequent biofilm formation. Our findings provide the first example of a microbial secondary metabolite promoting biofilm formation via a physicochemical approach, highlighting the significance of secondary metabolism in mediating critical processes related to the development of dental caries.

## Introduction

Dental caries (tooth decay) has been recognized as one of the most common bacterial infections and costly chronic conditions afflicting humans^1,2^. In most industrialized countries, 60–90% of children and the vast majority of adults are affected by dental caries. Annually, the global economic burden of treating tooth decay amounts to billions of dollars^3–5^. The development of dental caries is a complex process that mainly depends on the presence of microbial biofilms on the tooth surfaces (known as dental plaque)^2,6,7^. In natural settings, establishing and persisting in the human oral cavity presents significant challenges to the oral microbiota because of the plethora of immune and nonimmune defenses that the host brings to bear against the organisms. As an effective strategy for surviving in the oral cavity, the formation of biofilms not only physically protects microorganisms from environmental challenges, but also provides a microenvironment for accumulating acidic metabolites that demineralize enamel^6–8^.

As a natural and primary inhabitant of the human oral cavity, more specifically, dental plaque, *Streptococcus mutans* has long been acknowledged as the major etiological agent that is most closely associated with the initiation of dental caries. This is attributed to its ability to 1) form biofilms; 2) produce organic acids such as lactic acid; and 3) grow in low pH environments. *S. mutans* possesses mechanisms of adhering to teeth that are coated with salivary pellicles, building and developing biofilms, and interacting chemically and physically with various members of the oral microbes to influence the composition and pathogenic potential of mature biofilm communities^7–10^. The macromolecules of *S. mutans* for biofilm formation and development have been extensively investigated, including many surface associated proteins^10,11^, such as glycosyltransferases (Gtfs), glucan-binding proteins (Gbps), functional amyloids, and envelope-associated proteins. In contrast, except for a few quorum sensing signals, such as autoinducer-2 (AI-2) and competence stimulating peptides (CSPs)^10^, the role of small-molecule secondary metabolites in biofilm formation of *S. mutans* remains largely unexplored. Given the significance of *S. mutans* biofilms in dental caries initiation and development, exploration of any new molecular mechanisms underlying *S. mutans* biofilm formation could facilitate further study of approaches to inhibit *S. mutans*-induced dental caries.

Polyketides/non-ribosomal peptides, known as major families of specialized secondary metabolites, have been noted for their structural and functional diversities. These metabolites are used by microbes to access information about both intracellular physiological status and extracellular environment, and have been shown to be critical in controlling complex processes, such as virulence, stress responses, defense, and biofilm formation^12,13^. Recently, genomic analysis of various *S. mutans* isolates demonstrated that these facultative anaerobes collectively harbored a plethora of biosynthetic gene clusters (BGCs) for polyketide/non-ribosomal peptide biosynthesis^14–16^. Some of the associated products were also reported, including mutanobactin with an inhibitory activity against the morphological transition of *Candida albicans*^17,18^, mutanocyclin that suppressed the infiltration of leukocytes^19^, and reutericyclin that inhibited the growth of oral commensal bacteria^20^ (Fig. 1a). Collectively, these studies raise the question if *S. mutans* can employ these unexplored BGCs to promote other critical physiological processes, such as adhesion and biofilm formation, during the lifetime of these oral pathogens.

**Fig. 1.**
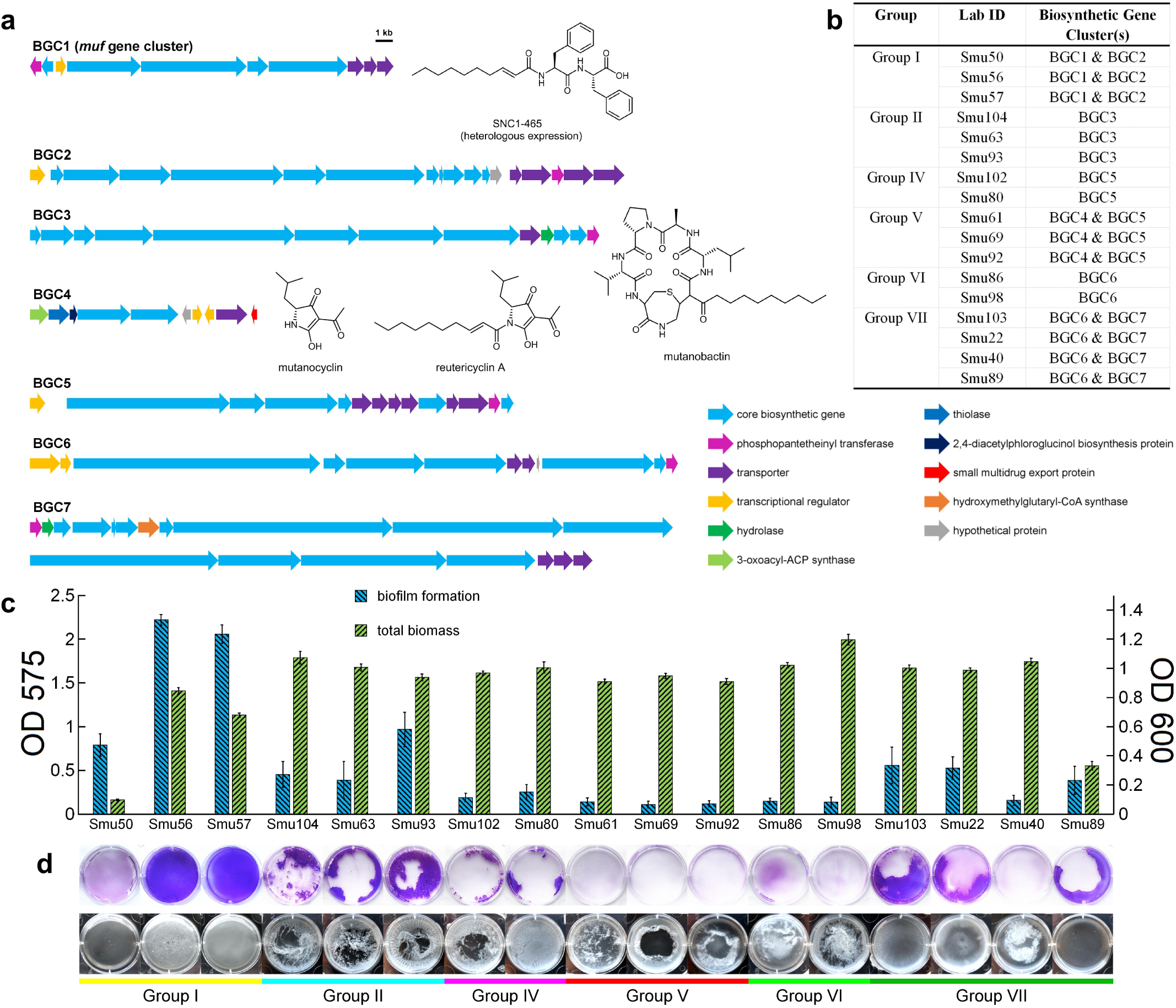
Biofilm formation by *S. mutans* isolates with diverse biosynthetic gene clusters (BGCs). **a**, Organizations of seven natural product BGCs, which are harbored in 17 selected *S. mutans* clinical strains. The structures of SNC1-465, mutanobactin, and mutanocyclin and reutericyclin A are shown, whose biosynthesis is linked with BGC1, BGC3, and BGC4, respectively. **b**, Distribution of BGCs amongst 17 *S. mutans* strains. **c**, Quantifications of biofilm formation and total biomass formation by 17 *S. mutans* strains in semi-defined biofilm medium by crystal violet biofilm assay (OD 575) and optical density measurement (OD 600), respectively. Data are shown as mean ± s.d. (*n* = 3). **d**, Images of biofilms formed by 17 *S. mutans* strains before (bottom panel) and after (top panel) crystal violet staining.

In previous studies, we have collected 57 *S. mutans* clinical isolates from individuals of known dental caries status worldwide^21^; these isolates displayed high degrees of genetic diversity^21^ and phenotypic heterogeneity^22^. In the present study, based on varying biofilm-forming capability, we identify a hybrid polyketide/non-ribosomal peptide BGC, *muf*, as a positive determinant for *S. mutans*-associated biofilm formation. Further study of biofilm-promoting mechanisms underlying *muf* at a molecular level is conducted, revealing the production of a novel lipopeptide mutanofactin-697 and its mode of action. This study provides the first example of dental biofilm formation directly affected by a microbial secondary metabolite through a rare physicochemical mechanism.

## Results

### Identification of the *muf* BGC affecting biofilm production in *S. mutans*

Based on antiSMASH analysis of secondary metabolite BGCs, we preliminarily classified 57 *S. mutans* strains that we previously collected into eight groups (I to VIII) depending on the presence of BGCs for polyketide/non-ribosomal peptide biosynthesis (Fig. 1a,b). It is notable that many of these BGCs are mutually exclusive in various *S. mutans* isolates, suggesting evolutionarily distinct origins of these BGCs and likely diverse biological functions. Furthermore, we selected 17 representative strains and assayed their capacities of developing biofilm using an established method^22^. Interestingly, strains from Group I, in particular Smu56 and Smu57, outperformed all other strains in forming stable biofilms attached to the bottom of six-well plates when glucose was used as a carbon source (Fig. 1c,d).

Group I strains harbor both BGC1 and BGC2, which were confirmed to be expressed in the semi-defined biofilm medium (BM) based on the transcriptional analysis. To probe possible roles of these two BGCs in biofilm formation, gene knockout was performed in Smu56 and Smu57 for these two BGCs, respectively. The disruption of BGC1 (named *muf* thereafter), but not BGC2, led to the formation of abnormal biofilms with a loose and sponge-like architecture (Fig. 2a), while the biomass accumulation was not affected in either mutant. In addition, the biofilm of Δ*muf* mutant was observed to partly float in the medium, loosely attach to the bottom of six-well plates, and could be easily removed with gentle shaking (Fig. 2b). To further confirm the observed phenotype was not limited to the polystyrene surface of six-well plates, we conducted a similar assay, but employed the acrylic resin artificial teeth, which are widely used in oral rehabilitation and have been used as an alternative surface to interrogate the adhesion of streptococci and biofilm formation^23^. Consistently, the biofilm formed by the Δ*muf* mutant on the artificial tooth surface was significantly diminished, as compared to that of the wild-type strain (Fig. 2c), supporting the critical role of *muf* in biofilm formation on various surfaces.

**Fig. 2.**
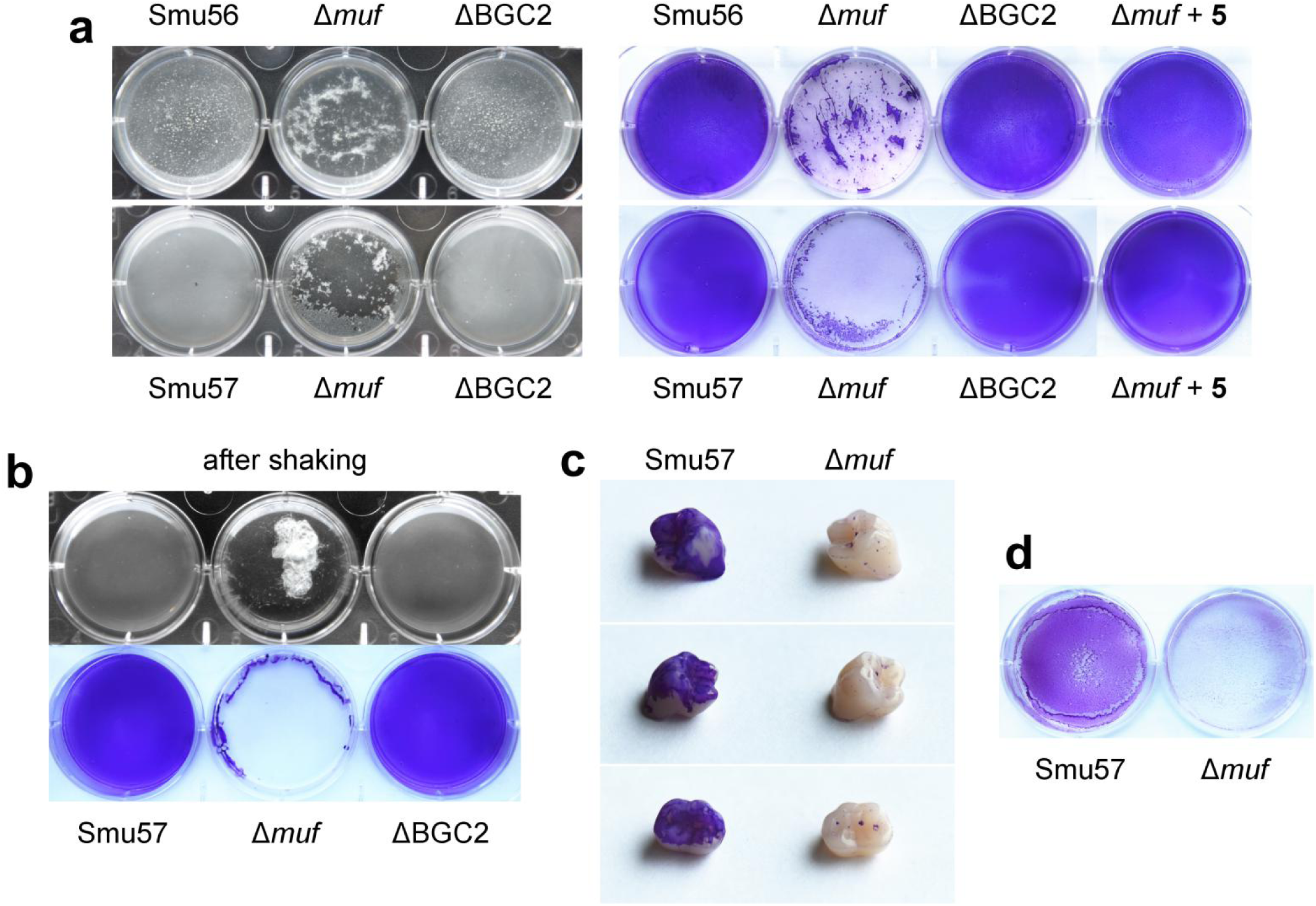
Biofilm formations by wild-type *S. mutans* strains and mutants. **a**, Images of biofilms formed by Smu56 and Smu57, their Δ*muf* mutants and ΔBGC2 mutants, and Δ*muf* mutants with complementation of compound **5**, attached to the bottom of polystyrene microtiter plates before (left panel) and after (right panel) crystal violet staining. **b**, The biofilm formed by Δ*muf* mutant showed abnormal architecture, and floated and twined together with gentle shaking. **c**, Artificial acryl teeth were incubated with Smu57 or Δ*muf* mutant, then the biofilm biomass developed was stained by crystal violet. **d**, The early stage biofilms (6-h-old) by Smu57 and Δ*muf* mutant were measured to evaluate the adhesion ability of these strains.

To probe whether additional gene(s) besides the *muf* BGC could synergistically contribute to the unique biofilm-forming capability observed in Group I strains, we constructed a Venn diagram which depicted all the shared and unique orthologous genes among these 57 *S. mutans* strains, and subsequently deleted all the unique genes (or gene clusters) harbored in Group I strains. The biofilm assays showed that the deletion of any of these unique genes (or gene clusters), but not *muf*, in Smu57 did not obviously impact biofilm formation. Collectively, these results suggested that the Group I strains-associated strong biofilm-forming capability depended solely on the presence of the *muf* BGC.

### *Muf* facilitates bacterial adhesion via increasing cell surface hydrophobicity

Given that surface adhesion and low pH resistance (aciduricity) are two crucial characteristics of *S. mutans* associated with biofilm formation, we next investigated the effect of the presence of *muf* on these two properties^7,10,24,25^. There was no observable difference in the acid tolerance of Smu57 between the wild-type and Δ*muf* strains, indicating that *muf* did not affect aciduricity of *S. mutans*. On the other hand, we observed a significantly decreased surface adhesion during the onset of the biofilm formation^26^ (Fig. 2d), suggesting that *muf* promoted biofilm formation of Group I strains by facilitating initial surface adhesion. In addition, using SYTO 9 (green) and propidium iodide (PI; red) for staining live and dead cells, respectively, a closer examination of the biofilms by confocal laser scanning microscopy (CLSM) revealed that the deletion of *muf* had no deteriorating impact on the cell viability, but caused significant dispersion of cells within the biofilm matrix.

Many studies have demonstrated that the bacterial cell surface hydrophobicity (CSH) plays an important role in promoting surface adhesion^24,27–29^. We thus measured and calculated the surface thermodynamic properties of different *S. mutans* wild-type strains and mutants using the well-established Sessile drop contact angle method. The water contact angle (*θ*_W_) has a predictive value for bacterial hydrophobicity, with a higher value correlated with a more hydrophobic bacterial surface^29,30^. The measurement demonstrated that the Group I strains had more hydrophobic surfaces than those of strains from any other groups; and three Δ*muf* mutants displayed *θ*W values decreasing from 62.45° ~ 68.64° to 44.85° ~ 46.09° (Fig. 3a,b), suggesting that this gene cluster is related to the bacterial CSH.

**Fig. 3.**
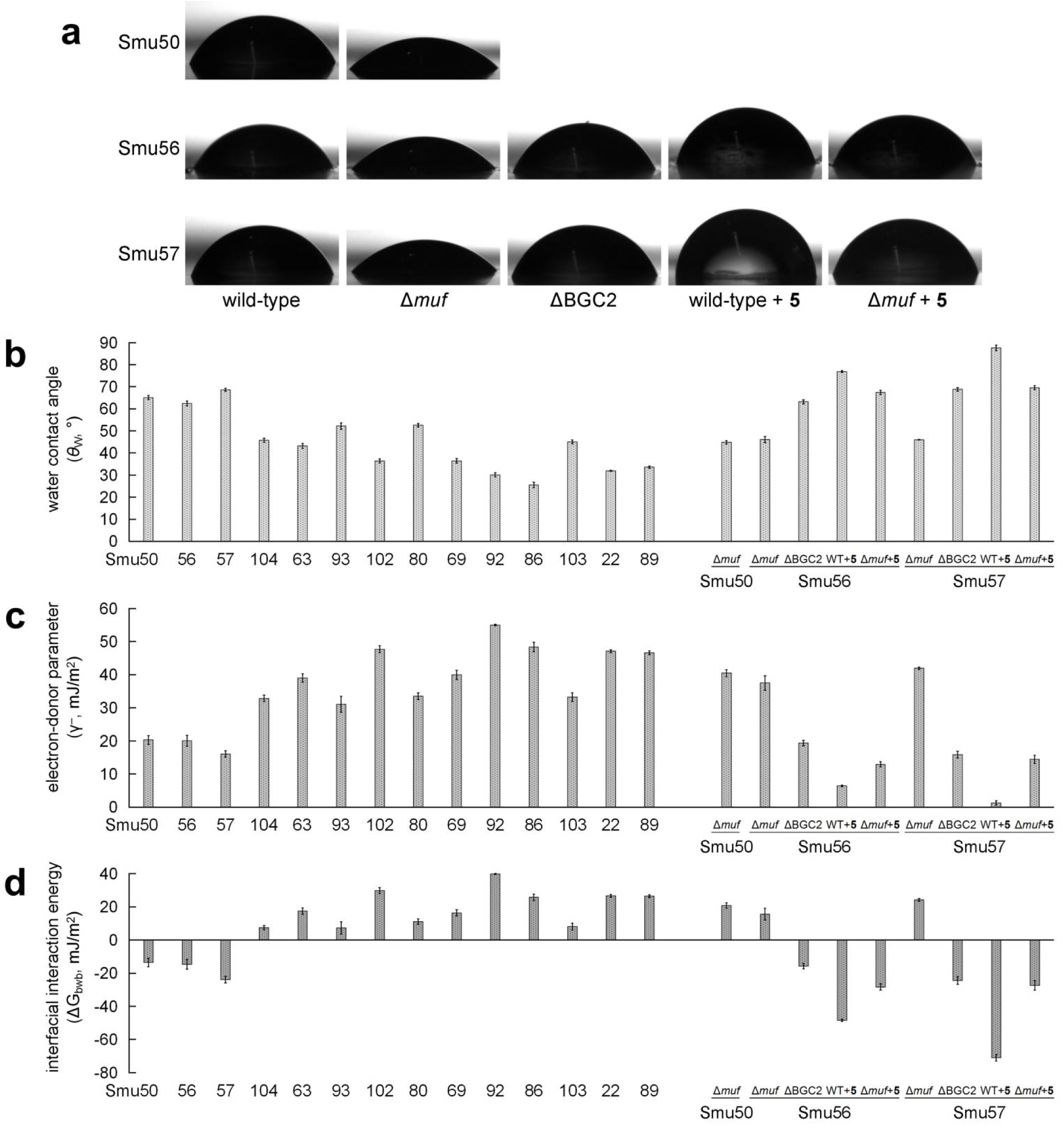
Surface physicochemical properties of *S. mutans* strains. **a**, Images of water contact angles which were measured by placing a drop of water onto a lawn of bacterial cells. **b**–**d**, Determination of the surface physicochemical parameters of different wild-type *S. mutans* strains and their mutants under various conditions using the contact angle method: water contact angle (*θ*W) (**b**), electron-donor parameter of the surface tension (γ^−^) (**c**), and interfacial interaction energy (ΔG_bwb_) (**d**). Data are shown as mean ± s.d. (*n* = 3).

We next included formamide and diiodomethane as the probe liquids for contact angle measurement (*θ*_FO_ and *θ*_DIM_), and calculated the cell surface tension parameters (γ^LW^, γ^−^ and γ^+^) and interfacial interaction energy (ΔG_bwb_) of *S. mutans* strains, to further estimate CSH. The polar electron-donor (γ^−^) parameter of a bacterial surface has been reviewed as a semiquantitative parameter in predicting bacterial CSH, where decreasing numerical value corresponds to increasing hydrophobicity^31,32^. All three *muf*^+^ strains had the lowest γ^−^ values (16.10 ~ 20.30 mJ/m^2^) among all tested wild-type strains (31.09 ~ 55.00 mJ/m^2^ for others). Consistently, all the Δ*muf* mutants exhibited significantly increased γ^−^ values (37.51 ~ 41.95 mJ/m^2^), compared to those of their wild-type counterparts (Fig. 3c). Based on the surface tension parameters, we then calculated the surface free energy of aggregation between bacterial cells in water (ΔG_bwb_), which is a quantitative measurement of CSH^31,32^. A negative number of ΔG_bwb_ indicates that the interaction between two cells is stronger than the interaction of each cell with water, and thus bacterial cells are apt to aggregate and are considered hydrophobic. Conversely, the cells are considered hydrophilic when ΔG_bwb_ is positive. Again, three *muf*^+^ strains differed from other group strains with negative values of ΔG_bwb_, demonstrating the unique hydrophobic property of the Group I strains in contrast to hydrophilicity of any *muf*^−^ strains (Fig. 3d). In addition, Δ*muf* switched the Group I strains from hydrophobic to hydrophilic, being consistent with the observed water contact angle measurements and the critical role of *muf* in increasing CSH (Fig. 3d). We thus propose that the presence of *muf* leads to a significant increase in cell surface hydrophobicity of *S. mutans*, which promotes the initial surface adhesion and the subsequent biofilm formation and maturation.

### Discovery of the *muf* products

Next, we turned to identifying the products of the *muf* BGC. We prepared the fermentation extracts of the wild-type Smu57 and its Δ*muf* mutant, compared their metabolite profiles using liquid chromatography–high resolution mass spectrometry (LC–HRMS), and succeeded in detecting five molecules with *m*/*z* of 459.2281 [*M* + H]^+^, 542.2121 [*M* + H]^+^, 540.2317 [*M* + H]^+^, 608.2216 [*M* + H]^+^ and 698.2537 [*M* + H]^+^ that were present only in the culture extracts of the wild-type Smu57 (Fig. 4). We named these five compounds as mutanofactin-458 (**1**), mutanofactin-541 (**2**), mutanofactin-539 (**3**), mutanofactin-607 (**4**) and mutanofactin-697 (**5**), respectively (Fig. 4). In addition, these compounds were also detected in the culture of other Group I strains, including Smu50 and Smu56, but not from the selected strains of other groups, demonstrating a strong correlation of these metabolites with *muf*. Among these five metabolites, mutanofactin-697 (**5**) was produced as the major product.

**Fig. 4.**
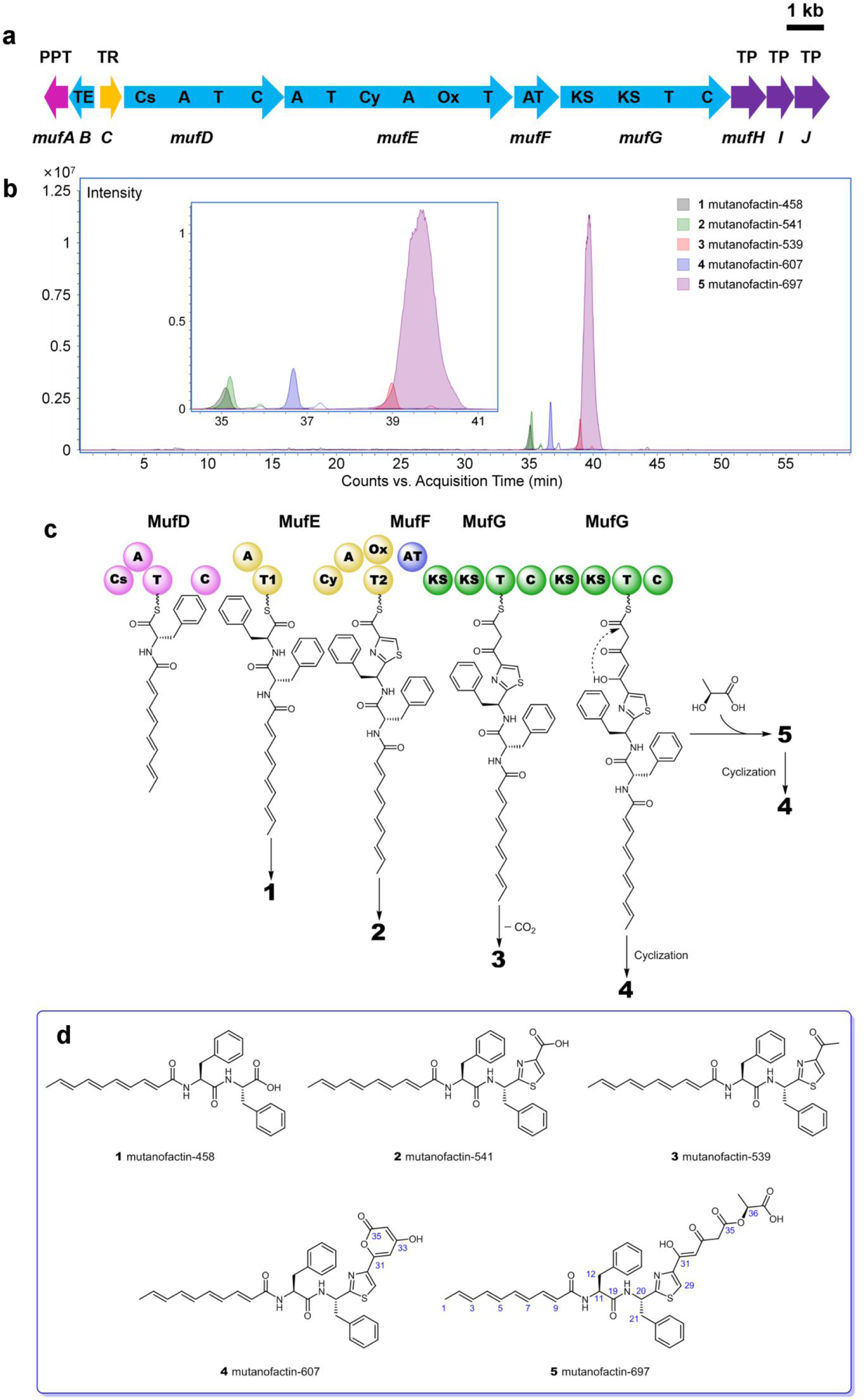
Biosynthesis and chemical structures of mutanofactins. **a**, Organization of the mutanofactin (*muf*) biosynthetic gene cluster. Genes encoding nonribosomal peptide synthetase (NRPS), polyketide synthase (PKS), and thioesterase (TE) are shown in blue; the 4′-phosphopantetheinyl transferase (PPT) gene *mufA* is shown in pink; the *muf* pathway transcriptional regulator (TR) gene *mufC* is shown in yellow; and the ABC transporter (TP) genes are shown in purple. A, adenylation; AT, acyltransferase; C, condensation; Cs, starter condensation; Cy, cyclization; KS, ketosynthase; Ox, oxidase; T, thiolation sequence of acyl-or peptidyl-carrier proteins. **b**, LC–HRMS extracted ion chromatogram traces of the metabolic extract of Smu57. EIC+ = 459.23 ± 0.01, 542.21 ± 0.01, 540.23 ± 0.01, 608.22 ± 0.01, and 698.25 ± 0.01, which correspond to compounds **1**, **2**, **3**, **4**, and **5**, respectively. **c**, Proposed biosynthetic pathway for mutanofactins. **d**, Chemical structures of mutanofactins.

From a 20 L fermentation culture of Smu57, 4 mg of **5** was obtained, after organic solvent extraction, followed by multiple rounds of reversed-phase liquid chromatography purification. Compound **5** was obtained as a white and amorphous powder with a molecular formula of C_38_H_39_N_3_O_8_S according to its HRESIMS data, suggesting 21 indices of hydrogen deficiency. Subsequently, a series of 1D (^1^H and ^13^C) and 2D (COSY, HSQC, and HMBC) NMR spectra of **5** was acquired, allowing assignment of all the carbons and protons of this new compound (Fig. 4d). Briefly, 5 was elucidated as a unique linear hybrid polyketide/non-ribosomal peptide hybrid metabolite that contains a (2*E*,4*E*,6*E*,8*E*)-deca-2,4,6,8-tetraenoyl tripeptide fragment, a linear triketide fragment, and a lactic acid moiety. The long-chain unsaturated fatty acyl moiety was proposed by proton-proton coupling constants of H-1 to H-9, and the COSY correlations between H-1/H-3 and H-2, H-3/H-5 and H-4, H-5/H-7 and H-6, and H-7/H-9 and H-8. The tripeptide fragment that contains two Phe residues and a Cys derived thiazole moiety was indicated by COSY correlations between 10-NH-/H-12a/b and H-11, and 19-NH-/H-21a/b and H-20, following with many HMBC cross-peaks. A linear triketide fragment was assigned in **5**, which consists of the carbons from C-31 to C-35. The remaining C_3_H_6_O_3_ was assigned as a lactic acid moiety, which was connected to the linear triketide acid precursor via an ester bond. This assignment was verified by 3*J* HMBC from H-36 to the ester carbonyl carbon at *δ*C 167.1 (C-35). The configuration of lactic acid was assigned to be l-after chemical degradation followed by Marfey’s analysis^33^. The chemical structure of **5** was further confirmed by multiple-stage MS*^n^* data from an ESI-LTQ Orbitrap XL MS spectrometer.

Additionally, the chemical structure of a minor metabolite, mutanofactin-607 (**4**), was determined in the same manner based on the 1D and 2D NMR data. **4** (molecular formula: C_35_H_33_N_3_O_5_S) shares the same chemical skeleton from C-1 to C-30 as that of **5**, and contains a terminal triketide lactone moiety that consists of the carbons from C-31 to C-35. Partial conversion of purified **5** to **4** was observed, suggesting that **4** could be a degradation product of **5**. The putative structures of additional minor metabolites, including **1**, **2**, and **3**, were further proposed based on a comparative analysis of their HR-MS/MS fragmentation data with those of **4** and **5** (Fig. 4d). The purified compound **1** was subjected to chemical degradation followed by Marfey’s analysis, which revealed the configuration of both Phe to be l-. This result suggested that **5** is most likely derived from two l-Phe residues, which is also consistent with the bioinformatic prediction of substrate specificity of *muf* pathway biosynthetic enzymes in the following discussion.

### Proposed biosynthetic pathway for mutanofactins

The identified structures of mutanofactins, together with the bioinformatic analysis of the *muf* gene cluster, allowed us to propose a putative enzymatic pathway for mutanofactin biosynthesis (Fig. 4). The linear scaffold of mutanofactins is presumably generated through a hybrid non-ribosomal peptide synthetase (NRPS)–polyketide synthase (PKS) assembly line-based mechanism utilizing two multi-domain NRPSs (MufD and MufE) and *trans*-AT PKSs (MufF and MufG). Specifically, MufD, with domains organized into Cs-A-T-C (Cs, starter condensation; A, adenylation; T, thiolation; C, condensation) is proposed to initiate chain assembly. The A domain is predicted to activate l-Phe, based on the known signature residues of binding-pocket for the Phe-specific A domains^34,35^. The Cs domain that shows homology to typical Cs domains involved in lipoinitiation is proposed to accept an acetate-derived (2*E*,4*E*,6*E*,8*E*)-deca-2,4,6,8-tetraenoyl-CoA as the substrate and acylate the amino substituent of the loaded l-Phe^36^. MufE is a dimodule NRPS with domains organized into A1-T1-Cy-A2-Ox-T2 (Cy, cyclization; Ox, oxidase). The A1 domain is predicted to also activate l-Phe, which is then condensed with *N*-acyl-l-Phe via an amide bond catalyzed by the terminal C domain of MufD. The resulting MufE-T1 tethered biosynthetic intermediate could be spontaneously hydrolyzed to produce the minor product **1** (Fig. 4c). The second module of MufE (Cy-A2-Ox-T2) shows homology to typical thiazole-forming modules, such as EpoB and MtaC in the biosynthesis of epothilone and myxothiazol, respectively^37,38^, and is thus predicted to extend the assembly line via incorporating an l-Cys-derived thiazole moiety. The utilization of Cys as a building monomer was supported by l-[1-^13^C]Cys feeding assays; and Cys was further demonstrated to be a limiting substrate for **5** production in various culture media (Fig. 5a,b). The resulting MufE-T2 tethered intermediate is then recognized by the downstream PKSs for further processing in generating the final product **5**; alternatively, a spontaneous hydrolysis in this step could yield the minor product **2**. The PKS MufG has domains organized into KS-KS-T-C (KS, ketosynthase) and is predicted to partner with MufF, a *trans*-acting acyltransferase (AT) that is malonate specific (signature motif: GAFH)^39–42^. MufF catalyzes two consecutive decarboxylative condensations using malonate extender units, presumably catalyzed by the two adjacent KS domains, respectively, followed by the release of the product from the assembly line by l-lactic acid via an ester bond formation promoted by the terminal C domain of MufG. It is notable that while a terminal condensation-like domain has recently received attention as a chain releasing mechanism in PKS/NPRSs^43–46^, particularly in fungal NRPSs, the terminal C domain promoted esterification remains rare, and no report has been associated with l-lactic acid, a major glycolytic product of *S. mutans*. The minor products **3** and **4** are plausible shunt products of PKS upon hydrolysis and decarboxylation for **3**, and synchronous cyclization and release for **4**; albeit **4** could also be formed from **5** after intramolecular cyclization and release of lactate.

**Fig. 5.**
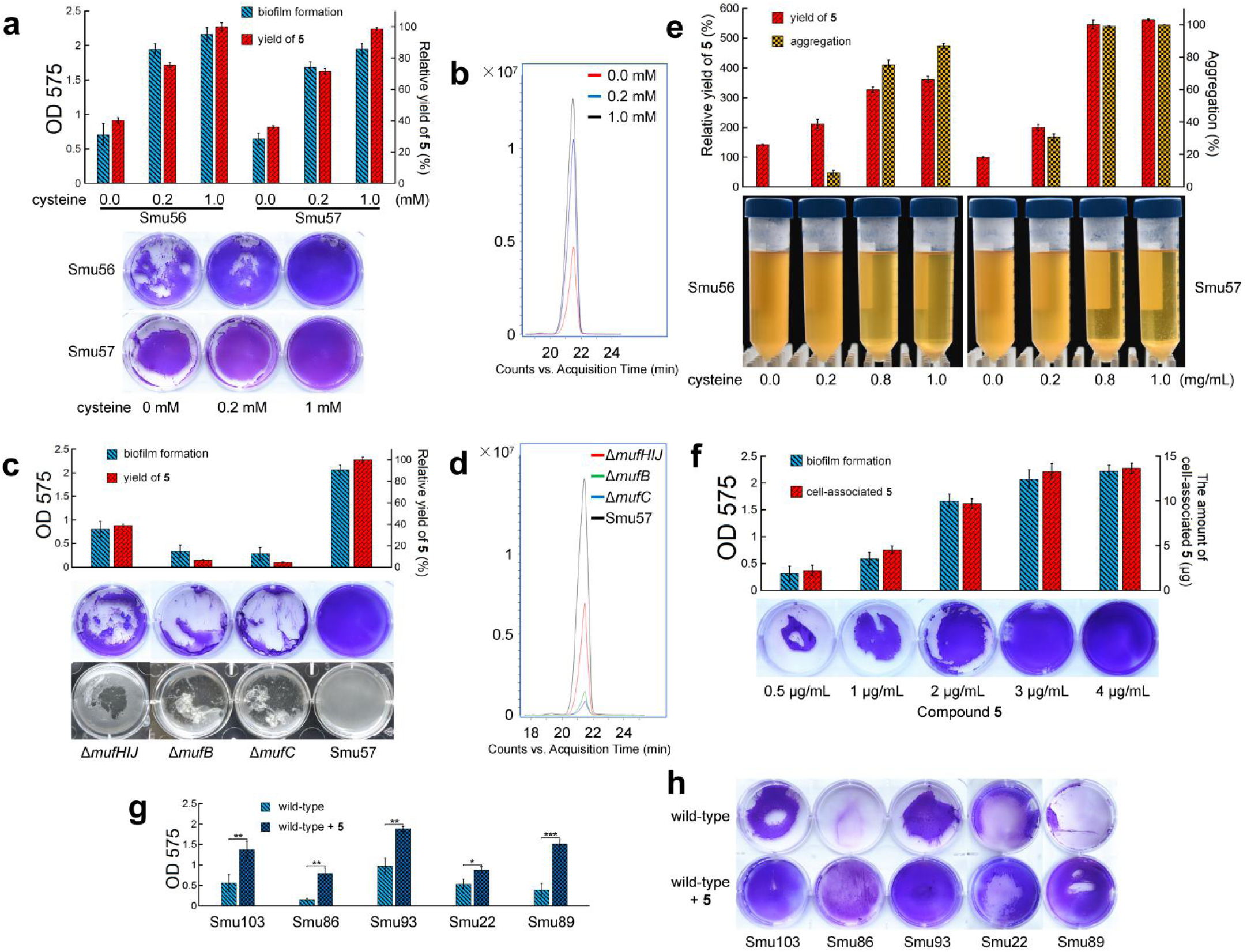
Effect of compound 5 on biofilm formation and cell self-aggregation. **a**, Quantification of biofilm formation and **5** production of Smu57 and its Δ*mufB*, Δ*mufC*, and Δ*mufHIJ* mutants. (upper panel) Mean value ± s.d. (*n* = 3); (bottom panel) Images of typical biofilms formed by different strains. **b**, LC–HRMS extracted ion chromatogram traces of the metabolic extracts of Smu57 and its three mutants. EIC+ = 698.25 ± 0.01, which corresponds to **5**. **c**, Quantification of biofilm formation and **5** production of Smu57 in biofilm media supplemented with Cys of different concentrations. (upper panel) Mean value ± s.d. (*n* = 3); (bottom panel) Images of typical biofilms formed by Smu57 under different culture conditions. **d**, LC–HRMS extracted ion chromatogram traces of the metabolic extracts of Smu57 under different culture conditions. EIC+ = 698.25 ± 0.01, which corresponds to **5**. **e**, Quantification of cell self-aggregation and **5** production of Smu56 and Smu57 under different culture conditions. (upper panel) Mean value ± s.d. (*n* = 3); (bottom panel) Images of Smu56 and Smu57 cultures in BHI media supplemented with Cys of different concentrations. **f**, Effect of increased concentration of **5** on biofilm-forming ability of Smu57 Δ*muf* in a **5**-complementation assay. In the assay, the amount of cell-associated **5** was measured. (upper panel) Mean value ± s.d. (*n* = 3). **g**,**h**, A comparison of biofilm formation of five wild-type *muf*^−^ strains with and without **5** complementation. (**g**) Quantification of biofilm formation. Data are shown as mean ± s.d. (*n* = 3). \**P* < 0.05; \*\**P* < 0.01; \*\*\**P* < 0.001; unpaired student *t*-test. (**h**) Images of typical biofilms formed by five wild-type *muf*^−^ strains with and without **5** complementation. **a**,**c**,**f**,**g**, Crystal violet biofilm assay (OD 575) was employed for quantifying biofilm formation.

In addition to the core biosynthetic NRPS/PKSs, the *muf* BGC is also predicted to encode a 4′-phosphopantetheinyl transferase (PPT, MufA) for post-translational modification of apo-T domains to the pantetheinylated active forms, a type II thioesterase (TE, MufB), a transcriptional regulator (TR, MufC), and three ATP-binding cassette (ABC) transporters (MufH, I, and J) (Fig. 4a). Type II TEs are known to enhance assembly line product formation by removing aberrant residues blocking the megasynthases, or participating in biosynthetic intermediate or product release^47^. Consistent with the predicted editing role of MufB, the Δ*mufB* mutant of Smu57 significantly decreased the production of **5** (Fig. 5c,d). Similarly, disruption of *mufC*, which encodes a predicted type II TetR family regulator^48^, also significantly decreased the production of **5**, suggesting the function of MufC as a positive regulator (Fig. 5c,d). In addition, simultaneous disruption of three transporter genes resulted in a moderate yield loss of **5**, implying the role of MufH, I, and J in exporting **5** extracellularly (Fig. 5c,d).

### Direct effect of mutanofactin on biofilm formation

The genetic manipulation of the *muf* BGC yielded a few Smu57 mutants with decreased production of **5**. Accordingly, we observed the disrupted ability of these mutants in biofilm formation, correlated with the yield of **5** (Fig. 5c,d). Similar correlations of the biofilm-forming ability (Fig. 5a,b), as well as spontaneous self-aggregation of bacterial cells (Fig. 5e), with the yield of **5** in multiple assays were also observed by varying the concentration of the limiting substrate Cys, suggesting that **5** may influence the biofilm formation and bacterial aggregation in a concentration-dependent manner.

We next performed the chemical complementation assay of **5** to directly probe its effect on biofilm formation. The addition of the purified compound **5** to the Δ*muf* mutant cultures successfully restored the biofilm formation in a dose-dependent way (Fig. 2a, and Fig. 5f). In addition, **5** induced macroscopically visible aggregation and classical biofilm ring formation of Δ*muf* mutant^49^. Furthermore, the addition of **5** to the wild-type or the Δ*muf* mutants of Smu56 and Smu57 significantly decreased the polar electron-donor (γ^−^) parameter of bacterial surface (Fig. 3c) and the surface free energy of aggregation (ΔG_bwb_) (Fig. 3d), demonstrating the role of **5** in increasing cell surface hydrophobicity of *S. mutans*.

To investigate broader biological impacts of **5**, we further performed the chemical complementation assay of **5** with five selected *muf*^−^ wild-type *S. mutans* strains. We observed that **5** promoted biofilm formation in all tested samples (Fig. 5g,h), accompanying the switch of these strains' surface properties from hydrophilic to hydrophobic (Fig. 3), demonstrating that *S. mutans* strains lacking the *muf* BGC could utilize extracellular **5** to increase their own CSH and improve biofilm formation. These results suggest that *muf*^−^ *S. mutans* strains may hijack **5** produced and secreted by neighboring *muf*^+^ strains in forming biofilms, which was supported by the cocultivation assays of Smu56 with six *muf*^−^ strains. It is notable that the exogenous **5** was found to be mainly cell-associated in chemical complementation assays with an estimated maximum of ~10^7^ molecules per cell, suggesting a possible mode of action underlying **5** in direct binding of bacterial cells to affect CSH due to the hydrophobicity of **5**.

To further probe the mechanism of **5** in increasing CSH, we examined the expression level of a few representative surface proteins that are known to be important for *S. mutans* cellular adhesion and biofilm formation^10,11^. The selected targets include GtfB, a glycosyltransferase synthesizing water-insoluble glucan rich in α-1,3-linkages; GbpA and GbpC, glucan-binding proteins that are important for adhesion; and SpaP, the I/II antigen interacting mainly with saliva agglutinins in the sucrose-independent adhesion and aggregation of *S. mutans*. The quantitative PCR analysis of the corresponding genes showed no difference in expression levels between the wild-type Smu57, the Δ*muf* mutant, and the **5**-complemented Δ*muf* mutant, arguing against these surface proteins-mediated CSH change.

## Discussion

Through extensive bioinformatic analysis, phenotypic screening, and gene deletions, we have identified a new polyketide/non-ribosomal peptide biosynthetic gene cluster (*muf*) that influences the biofilm formation of *S. mutans*, the major pathogen of dental caries. Combining both theoretical calculations and experimental measurements, we have shed light on the mechanism underlying the *muf*-promoted biofilm formation via facilitation of initial surface adhesion and cell–cell aggregation. The adhesion process is generally modelled by the Derjaguin–Landau–Verwey–Overbeek (DLVO) theory, in which initial adhesion consists of the balance between the attractive van der Waals force and electrostatic repulsion^27,31^. The van der Waals force is typically expressed as the Lifshitz–van der Waals component of surface tension (γ^LW^), and the electrostatic repulsive forces can be expressed in terms of the electron-acceptor parameter (γ^+^) and the electron-donor parameter (γ^−^) of Lewis acid–base component of surface tension. The surface free energy of aggregation between bacterial cells in water (ΔG_bwb_) can be calculated using these parameters obtained from cells and water, which is typically positive indicating hydrophilicity for most bacterial cells^32,50^. Consistently, all the wild-type strains in this study that lack *muf* display positive ΔG_bwb_ values and are dispersed in broth culture. The hydrophilicity of bacterial cells is known to be mainly influenced by the electron donor characteristic (γ^−^), which varies with the electron donating ability of surface components^31,32,50^. Notably, the presence of *muf* significantly decreases γ^−^ while minimally impacting γ^LW^ or γ^+^, being consistent with the predominant role of γ^−^ in predicting cell surface hydrophobicity (Fig. 3). Accordingly, ΔG_bwb_ decreases and switches from positive to negative due to the presence of *muf*, indicating increased cell hydrophobicity to promote cellular aggregation and likely surface adhesion.

To understand the role of *muf* at a molecular level, in the current study, we identified a novel group of small-molecule metabolites associated with *muf*, including lipopeptide mutanofactin-697 (**5**) that contains a unique molecular scaffold and displays as the major bioactive product (Fig. 4d). The biosynthesis of **5** can be readily predicted based on the *muf* BGC-encoded NRPS–PKS assembly line, which further supports the direct function of *muf* in producing mutanofactins. On the other hand, the structure of **5** embodies a couple of biosynthetic features which cannot be precisely predicted through *in silico* analysis only, such as the incorporation of a highly unsaturated fatty acyl moiety catalyzed by a starter condensation domain and l-lactic acid mediated off-loading mechanism promoted by a terminal condensation domain (Fig. 4c). Recently, a BGC homologous to *muf* from *S. mutans* NMT4863 has been subjected to heterologous expression and metabolite identification. In particular, the BGC was introduced into a model organism of *S. mutans* UA159 and activated via inserting two oppositely oriented constitutive promoters to replace the regulatory gene homologous to *mufC*. Unfortunately, only a shunt metabolite was identified which shares the same structural scaffold as mutanofactin-458 (**1**) except containing a different (2*E*)-decenoyl starting acyl moiety^19^ (Fig. 1a). This result supports our predicted structure of **1** and the associated biosynthetic function of *muf* genes, meanwhile showing limitation of employing a heterologous expression strategy for revealing native BGC products.

The discovery of mutanofactin-697 (**5**) and its biosynthetic pathway provides a unique opportunity to understand the role of **5** in biofilm formation through a quantitative analysis. **5** was found to bind to *S. mutans* cells, modify bacterial surface physicochemical properties (decreasing both γ^−^ and ΔG_bwb_ values), and promote self-aggregation of bacterial cells and biofilm formation in a dose-dependent manner. Particularly, **5** was active toward not only the native producers, but genomically-diverse *S. mutans* isolates lacking *muf*, indicating a general mode of action underlying **5**. It is notable that the BGC homologous to *muf* has also been detected in other streptococci species, such as *Streptococcus mitis* SK597 from the human mouth and *Streptococcus equinus* Sb10 from horse feces^19^, suggesting a potentially widespread activity of **5** in the microbial ecology of streptococci. Here, we propose the following mechanism for biofilm formation promoted by mutanofactin-697 (**5**), based on all the results obtained in this study. After being biosynthesized and secreted from a producing streptococcus, **5** binds to self and neighboring streptococci, increases bacterial cell surface hydrophobicity, promotes initial bacterial adhesion and subsequent biofilm formation and maturation. Although the exact mechanism of **5** in binding bacterial cells and increasing surface hydrophobicity remains unclear, we postulate that **5** nonspecifically binds to the cell surface and increases bacterial hydrophobicity physicochemically, considering the unique chemical structure of **5** and the high number of bound **5** per cell. However, we do not exclude the possibility that **5** induces signaling processes that change surface proteins and glucans left out in our transcription assays.

In summary, we discover a novel bacterial small molecule, mutanofactin-697, which directly contributes to a strong biofilm-forming capability of *S. mutans* clinical isolates. Further mode-of-action studies reveal that mutanofactin-697 binds to *S. mutans* cells nonspecifically, increases bacterial hydrophobicity, and promotes biofilm formation via an enhanced initial adhesion capability. We postulate that mutanofactin-697 may influence dental biofilm composition and architecture by binding to other plaque microorganisms in addition to *S. mutans*. Our discoveries thus highlight the significance of microbial secondary metabolism in mediating critical processes, such as biofilm formation in human oral pathogens, and enable further mechanistic interrogation of mutanofactin-related chemical regulatory processes in human oral ecology and streptococci-induced dental caries incidence and prevention.

## Acknowledgements

This work was financially supported by grants from the Chan Zuckerberg Biohub Investigator Program to W.Z., the Hong Kong Branch of Southern Marine Science and Engineering Guangdong Laboratory (Guangzhou) to W.Z. and P.-Y.Q. (SMSEGL20SC01), the China Ocean Mineral Resources Research and Development Association grant (COMRRDA17SC01) and National Key R&D Programmes of China (MOST19SC04) to P.-Y.Q., and the National Institutes of Health grant (R01 DE13239) to R.A.B. We thank J. G. Pelton (QB3, UC, Berkeley) for assistance with NMR measurements, A. T. Iavarone (QB3, UC, Berkeley) for assistance with mass spectrometry experiments, and D. Schichnes (The CNR Biological Imaging Facility) for assistance with microscope experiments.

## Author contributions

Z.-R.L. performed the experiments, Z.-R.L. and Y.D. analyzed the NMR data, Z.-R.L. and J.S. performed the gene analysis, Z.-R.L. and A.P. performed the contact angle measurements, L.Z. and R.A.B. assisted with *S. mutans* culturing and genetics, Z.-R.L. and W.Z. designed the study and wrote the manuscript, with input from R.M., P.-Y.Q., and R.A.B.

## Methods

### Detection of putative natural product biosynthetic gene clusters (BGCs)

Genome sequence data for 57 *S. mutans* clinical isolates have been deposited in GenBank^21^. Putative NRPS and/or PKS secondary metabolite BGCs were originally identified using antiSMASH version 5.0 with default settings^51^. Annotations were refined manually using CDsearch and BLASTp (Basic Local Alignment Search Tool) to identify conserved domains. The domain organisation of NRPS and/or PKS BGCs identified by antiSMASH was further analysed using the NRPS/PKS database^52^. The A domain substrate specificity for NRPS enzymes was predicted using NRPSpredictor2^53^, and NaPDoS was used to identify C and KS domains^54^.

### Bacterial strains, media, and growth conditions

17 *S. mutans* strains were selected from the 57 clinical collection. All strains were stored in 25% glycerol at –80 °C and freshly streaked on brain heart infusion (BHI) agar plates before each experiment. The routine culture of a *S. mutans* strain was inoculated from a single colony and grown in BHI medium (Difco™) at 37 °C in a 5% CO_2_ atmosphere. For biofilm experiments, strains were grown in semi-defined biofilm medium (BM) supplemented with 20 mM glucose^22^. The BM contained 58 mM K_2_HPO_4_, 15 mM KH_2_PO_4_, 10 mM (NH_4_)2SO_4_, 35 mM NaCl, 2 mM MgSO_2_**•**7H_2_O, and 0.2% (wt/vol) casamino acids, and was supplemented with filter-sterilized vitamins (0.04 mM nicotinic acid, 0.1 mM pyridoxine HCl, 0.01 mM pantothenic acid, 1 mM riboflavin, 0.3 mM thiamin HCl, and 0.05 mM d-biotin), and amino acids (4 mM l-glutamic acid, 1 mM l-arginine HCl, 1.3 mM l-cysteine HCl, and 0.1 mM l-tryptophan). In addition, different l-cysteine-deficient BM were prepared, with the final concentrations of 0 mM, 0.2 mM, and 1.0 mM.

### Biofilm assays in 6-well plate

Biofilm development was measured in a polystyrene 6-well cell culture plate (Costar 3516; Corning Inc., Corning, NY) as previously described^22^, with the following modifications. Overnight cultures were sub-cultured 1:50 into fresh 10 mL BHI medium and grown to mid-exponential phase (OD_600_ = 0.5). Then each culture was sub-cultured 1:70 into 7 mL BM medium in a 6-well cell culture plate, followed by incubation at 37 °C in a 5% CO_2_ atmosphere for 24 h without agitation. After 24 h, biofilms formed in 6-well plates were photographed immediately before staining. For staining, culture medium was removed by aspiration and wells were gently washed with 2 mL sterile deionized water. Subsequently, 1 mL of 0.1% crystal violet was applied to each well and incubated at room temperature for 15 min, followed by removing fluid by aspiration. Wells were washed twice with 2 mL sterile deionized water as before and allowed to air dry. The plates were photographed and the wells were destained with 2 mL of an acetone:ethanol solution (2:8) for 30 min at room temperature. The destaining procedure was repeated and the OD_575_ of the pooled destaining solution was measured. Using another copy of cultured biofilm sample, the total biomass (OD_600_) after 24 h incubation was measured after agitating and scraping all the attached bacterial cells.

### Biofilm assays on artificial tooth

Artificial teeth were applied for a static biofilm assay. Teeth were placed in 6-well plates. Each well of 6-well plates contained a 7 mL bacterial suspension of *S. mutans* in BM medium. After 24 h incubation at 37 °C, each tooth was washed twice with sterile deionized water to remove the nonattached cells, and the biofilm biomass was stained with crystal violet solution for 5 min at room temperature. The stained biofilm, which was formed on the teeth surfaces, was washed five times with sterile deionized water and then photographed.

### Evaluation of *S. mutans* adhesion

To evaluate *S. mutans* adhesion on microtiter dish, 6-h-old biofilms were evaluated^26^, which were prepared by the same method mentioned above.

### Transcriptional analysis of *muf* and BGC2

After culturing for 24 h in BM, the total RNA of Smu57 strain was isolated using RNeasy Mini Kit (QIAGEN) according to manufacturer’s protocol. To eliminate the DNA contamination, the total RNA was treated by RNase-free DNase Set (QIAGEN). The cDNA was prepared by using the First Strand cDNA Synthesis Kit following the manufacturer’s instructions (New England BioLabs). The same PCR program was applied to the total RNA sample after DNase treatment as negative controls.

### Construction of mutant strains

Standard DNA manipulation techniques were used to engineer plasmids and strains. Various BGC-deficient strains of *S. mutans*, including Δ*muf* mutants and ΔBGC2 mutants, were created using a PCR ligation mutagenesis approach^55^ to replace nearly all of the open reading frame (ORF) with a nonpolar kanamycin resistance marker (NpKm)^56^. For the selection of antibiotic-resistant colonies after genetic transformation, kanamycin (50 μg/mL for *E. coli* or 1 mg/mL for *S. mutans*) was added to the medium and agar plates when needed. Synthetic XIP (sXIP) (amino acid sequence, GLDWWSL), was synthesized and purified to 96% homogeneity by NeoBioScience (Cambridge, MA). The lyophilized sXIP was reconstituted with 99.7% dimethyl sulfoxide (DMSO) to a final concentration of 2 mM and was stored in 40 μl aliquots at –20 °C.

### Gene family analysis

Similarity among the protein sequences from seventeen strains were BLAST against each other with the threshold of 1e–5, and the orthologues were determined with OrthoMCL pipeline v2.0.9^57^ with the setting of “-I 1.5”. A total of 2521 orthologues were determined, and the Venn diagram was performed by the online tool (http://bioinformatics.psb.ugent.be/webtools/Venn/).

### Phylogenetic analysis

Protein sequences were aligned by ClustalW. Phylogenetic analysis was performed with MEGAX^58^. A Maximum Likelihood tree was constructed with 100 bootstrap replications, and the model of LG+Γ+I was selected as the best model.

### Acid tolerance assays

The effect of pH on the growth of the wild-type Smu57 and its Δ*muf* mutant was evaluated by assessing their growth on BHI agar plates with adjusted pH levels. Both the tested strains were grown in BHI medium (pH 7.0) overnight. 1 mL of overnight culture was transferred into 9 mL fresh medium, and incubation continued for 2 h at 37 °C. The cultures were gently sonicated for 15 s to disperse the chains of cells prior to serial dilution with 10 mM KPO_4_ buffer (pH 7.2). An aliquot (20 μL) of cell suspension from each strain was inoculated onto BHI agar plates with pH 4.6, pH 5.2, and pH 6.0. The plates were then incubated anaerobically at 37 °C for 48 h before assessing the acid sensitivity.

### Confocal laser scanning fluorescence microscopy

Confocal laser scanning microscopy (CLSM) was performed to observe different *S. mutans* biofilms. Strains were cultivated and grown in BM medium in a 6-well plate with coverslips. After 24 h, the coverslips were removed from the medium and were gently rinsed with a 0.85% saline solution. Coverslips were then stained with a LIVE/DEAD BacLight Bacterial Viability Kits (Molecular Probe, Eugene, OR, USA), following the manufacturer’s instructions. Viability of bacteria within the biofilms was determined: viable bacterial cells were stained with SYTO-9 (green), and bacterial cells with damaged membranes were stained with propidium iodide (PI; red). The stained biofilms were examined using a Zeiss LSM 880 FCS Confocal microscope with a 100x objective at both the green channel (488 nm excitation, green emission) and the red channel (543 nm excitation, red emission).

### Contact angle measurement

Strains were grown in 7 mL BM medium in 6-well cell culture plates. After 24 h of incubation, microbial biofilm was harvested by scraping using a 0.85% saline solution. After centrifugation at 4000 rpm for 10 min, the pellet was washed twice with saline solution and suspended in the same solution. Bacterial cells were counted, and bacteria suspension was filtered through a cellulose acetate membrane filter (0.45 μm) to deposit a uniform bacterial lawn of approximately 2 × 106 cells/mm^2^. Prior to measuring contact angles, the filters were air dried at room temperature for 30 min.

The contact angles were determined using a Ramé Hart 290-F1 automated goniometer by the sessile drop method. Droplets of three liquids (water, formamide, and diiodomethane) were applied to each surface. For contact angle measurements, a drop of 2 μL of the test liquid was dispensed on the surface of the filter. The contact angles were taken 2 s after drop deposition^59^.

### Surface tension components

The theory of surface tension components considers the surface tension to be composed of three separate components: Lifshitz-van der Waals component, which is nonpolar in nature, and electron donor/electron acceptor (Lewis acid/base) components, which are polar. The surface tension components are related to the contact angle through application of Young’s equation. For bacterial lawns, this is written as^31^

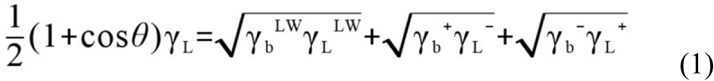

where γ_L_ is the total surface tension of the liquid, γ^LW^ is the Lifshitz-van der Waals surface tension component, γ^+^ is the electron acceptor component, γ^−^ is the electron donor component, and *θ* is the contact angle of the droplet on the bacteria lawn. The subscripts L and b stand for the liquid used for determining the contact angle (e.g., water) and the bacterial lawn, respectively. Through use of three different fluids with known surface tension components, the three unknown’s, γ_b_^LW^, γ_b_^+^, and γ_b_^−^, can be determined. The total surface tension of the bacteria lawn, γb, is then found as^31^

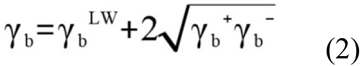

### Cell surface hydrophobicity

The bacterial cell surface hydrophobicity (CSH) can be determined via an analysis of the surface tension components. A hydrophobic surface is defined as one for which the free energy of hydrophobic interaction between two similar surfaces immersed in water is less than zero. For the bacterial lawn, this is written as^31^

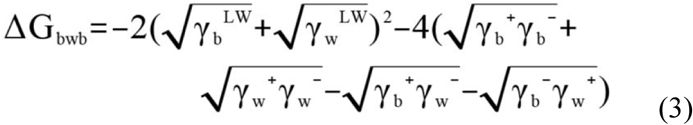

where the subscripts b and w stand for the bacterial lawn and water, respectively. Given eqs (1) and (3), the bacterial cell surface tension components and CSH can be determined through analysis of the contact angles.

### LC–HRMS-based analysis of *muf* pathway metabolites

For liquid chromatography–high resolution mass spectrometry (LC–HRMS)-based analysis, a single colony of the strain tested was picked for culture. After incubation, the culture broth was extracted with an equal volume of ethyl acetate (EtOAc) twice. The organic phase extract was separated by centrifugation (10 min, 4000 rpm), evaporated, and finally resuspended in 200 μL of methanol (MeOH). The prepared extract was subjected to LC–HRMS for analysis with injection volume of 10 μL.

LC–HRMS analysis was performed using an Agilent Technologies 6545 Accurate-Mass QTOF LC–MS instrument fitted with an Agilent Eclipse Plus C18 column (4.6 × 100 mm) by a gradient elution (A, H2O with 0.1% formic acid (FA); B, CH3CN with 0.1% FA: 2% B over 2 min, 2–100% B from 2 to 53 min, 100% B from 53 to 55 min, 100–2% B from 55 to 55.1 min, and 2% B from 55.1 to 60 min; flow rate, 0.5 mL/min). In addition, a 30-min elution condition was set for quick detection of target compounds: 2% B over 2 min, 2–100% B from 2 to 23 min, 100% B from 23 to 25 min, 100–2% B from 25 to 25.1 min, and 2% B from 25.1 to 30 min. The MS settings were as follows: positive ion mode, capillary voltage: 3500 V; nebulizer pressure: 40 psi; drying gas: 10 L/min; gas temperature: 320 °C; sheath gas flow: 11 L/min; sheath gas temperature: 350 °C; fragmentor voltage: 150 V; skimmer voltage: 65 V; and octopole RF: 750 V. LC–MS measured accurate mass spectra were recorded across the range 100–1700 *m*/*z*. All MS data were analyzed using Agilent MassHunter Qualitative Analysis software.

### Feeding experiments for biosynthetic pathway study

A single colony of the strain tested was picked for culture. In the next day, 100 μL of seed culture was inoculated in 10 mL of fresh BHI medium supplemented with 0.25 mg/mL l-[1-^13^C]cysteine, or 0.5 mg/mL [1-^13^C1]acetate, or 0.5 mg/mL [1,2-^13^C_2_]acetate (all the reagents were purchased from Cambridge Isotope Laboratories, Inc) in a 50 mL sterile Falcon tube, and incubated at 37 °C for 24 h. After incubation, each sample was treated as the method described above for LC–HRMS analysis.

### Fermentation and isolation of mutanofactin-607 (4) and mutanofactin-697 (5)

A single colony of the wild-type Smu57 was picked from a freshly streaked BHI agar plate and inoculated into 50 mL of BHI medium. After an overnight culture, 10 mL of culture was added to 2 L of fresh BHI medium supplemented with 1 mg/mL l-cysteine. In total, 40 L of BHI medium was cultured and bacterial cells were harvested by centrifugation (10 min, 4000 rpm). The collected bacterial cells were subjected to nonpolar extraction (MeOH and acetone, 5:1) twice. EtOAc was used to extract remaining compounds from the supernatant. The organic extracts were concentrated in vacuo, combined, and subjected to a flash chromatography system (Sepra C18 sorbent, 250 g). The crude extract was fractionated by elution with an MeOH–H2O gradient (50:50, 80:20, and 100:0) to yield three fractions (F01–F03). LC–MS analysis showed that the fraction F03 contained both **4** and **5**, which was further subjected to two rounds of HPLC purification with a semi-preparative C18 Phenomenex Luna column (5 μm, 250 mm × 10 mm inner diameter). Briefly, in the first round of HPLC separation, F03 was eluted with an CH_3_CN–H_2_O gradient mixture (A, CH3CN; B, H2O; 35% A over 2 min, and 35–75% A from 2 to 75 min) at a flow rate of 3.5 mL/min to obtain subfractions F0305 (*t*_R_ = 48–53 min) that was detected containing **4**, and F0307 (*t*_R_ = 56–60 min) that was detected containing **5**. F0305 and F0307 were purified through HPLC again, respectively, with an CH_3_CN–H_2_O gradient mixture (A, CH_3_CN; B, H_2_O; 30% A over 2 min, and 30–80% A from 2 to 80 min) at a flow rate of 3.5 mL/min to obtain **4** (*t*_R_ = 34–35 min; 2 mg) and **5** (*t*_R_ = 38–42 min; 40 mg). [We firstly purified 4 mg of **5** from a 20 L fermentation culture of Smu57, using BHI medium without supplementation of l-cysteine]

### NMR characterization of 4 and 5

^1^H, ^13^C, ^1^H–^1^H COSY, ^1^H–^13^C HSQC, and 1H–^13^C HMBC NMR spectra for **4** and **5** were acquired, respectively, on a Bruker Avance 900 NMR spectrometer (900 MHz for 1H and 225 MHz for ^13^C) equipped with a cryoprobe. For the NMR tests, both samples were dissolved in DMSO-*d*_*6*_ (Cambridge Isotope Laboratories, Inc) and loaded into 2.5 mm NMR tubes. Data were collected and reported as follows: chemical shift, integration multiplicity (s, singlet; d, doublet; t, triplet; m, multiplet), and coupling constant. Chemical shifts were reported using the DMSO-*d_6_* resonance as the internal standard for 1H-NMR DMSO-*d_6_*: δ = 2.50 ppm and ^13^C-NMR DMSO-*d_6_*: δ = 39.5 ppm.

### MS/MS and MS*^n^* analyses of mutanofactins

All of five mutanofactins (compounds **1** to **5**) were analyzed by LC–HRMS/MS, which was performed using the same LC condition described above with the target mass *m*/*z* 459.23 [*M* + H]^+^ and collision energy of 15 V around the retention time 35.1 ± 0.3 min for **1**; the target mass *m*/*z* 542.21 [*M* + H]^+^ and collision energy of 15 V around the retention time 35.2 ± 0.3 min for **2**; the target mass *m*/*z* 540.23 [*M* + H]^+^ and collision energy of 15 V around the retention time 38.9 ± 0.3 min for **3**; the target mass *m*/*z* 608.22 [*M* + H]^+^ and collision energy of 15 V around the retention time 36.7 ± 0.3 min for **4**; and the target mass *m*/*z* 698.25 [*M* + H]^+^ and collision energies of 15 V and 45 V around the retention time 39.8 ± 0.3 min for **5**. In addition, MS*n* fragmentation data of **5** was acquired using an ESI-LTQ Orbitrap XL (Thermo Fisher Scientific GmbH, Bremen, Germany) MS spectrometer.

### Marfey’s analysis

The absolute configuration(s) of phenylalanine(s) present in **1** were determined using Marfey’s methodology^60^. The acid hydrolysate of **1** was treated with Marfey’s reagent, 1-fluoro-2,4-dinitrophenyl-5-l-alanine amide (l-FDAA), and the resulting l-FDAA derivatives were analyzed by LC–HRMS. A modified Marfey's method (*O*-Marfey method)^33^ was employed for the determination of the configuration of lactic acid in **5**. Briefly, the hydrolysate of **5** was evaporated to dryness and resuspended in 1 mL of THF with sodium hydride (1 mg, 5mM) was added at room temperature. After 5 min, 200 μL of l-FDAA solution in THF (8 mg/mL, 6 mM) was added, and the reaction mixture was stirred under a nitrogen atmosphere for 5 min. A 30 μL aliquot of the reaction mixture was removed and quenched by adding 10 μL of 2 N HCl before LC–HRMS analysis. d-lactic acid and l-lactic acid were treated with l-FDAA in the same manner as the standards.

### Cell aggregation

Wild-type strains Smu56 and Smu57, and their Δ*muf* mutants were inoculated into 35 mL of BHI medium supplemented with 0.2 mg/mL, or 0.8 mg/mL, or 1 mg/mL l-cysteine. After 24 h incubation at 37 °C, the aggregation of each samples was determined by the turbidity variation of the bacterial suspension. The suspension was transferred into a cuvette, and the turbidity of suspension was determined at 600 nm by a spectrophotometer. The percentage aggregation was calculated as follows:

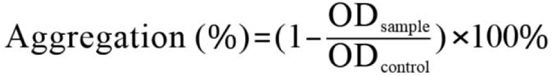

The OD_600_ value of BHI culture without supplemented l-cysteine was used as control.

### Treatment of *muf* mutant cells with 5

Smu57 Δ*muf* mutant was inoculated into 35 mL of BHI medium. After 24 h incubation at 37 °C, bacterial cells were harvested by centrifugation (10 min, 4000 rpm), and resuspended in 0.85% saline solution. Bacterial cells were washed twice with the same saline solution to remove soluble extracellular polymeric substance. After centrifugation again, the cell pellet was resuspended in **5** solution (2 μg/mL or 5 μg/mL). The suspensions were mixed in a shaker at 37 °C, 100 rpm for 2 h to achieve the adsorption equilibrium of **5**. The bacterial suspension in 0.85% saline solution with DMSO served as a control under which no **5** treatment was given.

### Compound complementation assay

For chemical complementation of Δ*muf* mutants, purified **5** was supplemented in each cultures at the time of inoculation with final concentrations of 0.5, 1.0, 2.0, 3.0, and 4.0 µg/mL. For chemical complementation of five selected wild-type *muf*^−^ strains, purified **5** was supplemented in each cultures at the time of inoculation with final concentration of 3.0 µg/mL. In each control group, the equivalent volume of DMSO was added. These controls showed no change in their biofilm formation

### Gene expression analysis of surface proteins

Quantitative PCR (q-PCR) was used to determine the expression levels of the *gtfB*, *gbpA*, *gbpC*, and *spaP* genes^10,11^. The protocols used for isolation, purification, and reverse transcription of total bacterial RNA into cDNA were described above. The amplifcation was performed using a StepOnePlus Real-Time PCR system with Fast SYBR Green Master Mix. 16S rRNA was used as an internal control.

### Co-culture experiments

Co-culture biofilm was prepared by inoculating the Smu56 and another *S. mutans* strain mixed bacterial suspension (in a 1:1 ratio) in 7 mL of BM. After 24 h of co-culture, crystal violet biofilm assay (OD_575_) was employed for quantifying biofilm formation, and colony-forming units (CFUs) of each strains were determined by counting the number of each strains (for Smu22, Smu98, and Smu93 that form distinguishing colonies) or colony PCR.

### Statistical analysis

All data were obtained from three independent assays. Data were expressed as mean ± s.d. for each group. The statistical significance of changes in different groups was evaluated by unpaired student *t*-test using GraphPad Prism software. The statistical significance at *P* < 0.05 was considered as significant.

